# Pollen thermotolerance of a widespread plant in response to climate warming: possible local adaptation of populations from different elevations

**DOI:** 10.1101/2023.06.22.546116

**Authors:** Karolína Hrubá, Paolo Biella, Jan Klečka

## Abstract

One of the most vulnerable phases in the plant life cycle is sexual reproduction, which depends on effective pollen transfer, but also on the thermotolerance of pollen grains. Pollen thermotolerance is temperature-dependent and may be reduced by increasing temperature associated with global warming. A growing body of research has focused on the effect of increased temperature on pollen thermotolerance in crops to understand the possible impact of temperature extremes on yield. Yet, little is known about the effects of temperature on pollen thermotolerance of wild plant species. To fill this gap, we selected *Lotus corniculatus* s.l. (Fabaceae), a species common to many European habitats and conducted laboratory experiments to test its pollen thermotolerance in response to artificial increase in temperature. To test for possible local adaptation of pollen thermal tolerance, we compared data from six lowland (389 - 451 m a.s.l.) and six highland (841 - 1030 m a.s.l.) populations. We observed pollen germination *in vitro* at 15, 25, 30, and 40°C. While lowland plants had stable germination rate at a wide range of temperatures between 15 and 30°C, with reduced germination rate observed only at extremely high temperatures (i.e., 40°C), the germination rate of highland plants was reduced already when the temperature reached 30°C, temperature commonly exceeded in the lowland during warm summers. This suggests that lowland populations of *L. corniculatus* may be locally adapted to higher temperature for pollen germination. On the other hand, pollen tube length decreased with increasing temperature in a similar way in lowland and highland plants. The overall average pollen germination rate significantly differed between lowland and highland populations, with highland populations displaying higher germination rate. On the other hand, the average pollen tube length was slightly smaller in highland populations. In conclusion, we found that pollen thermotolerance of *L. corniculatus* is reduced at high temperature and that the germination of pollen from plant populations growing at higher elevations is more sensitive to increased temperature, which suggests possible local adaptation of pollen thermotolerance.

## Introduction

Climate change, in particular increasing temperature, represents a major threat to biodiversity and is already leading to changes in the geographic distribution of many species of plants and animals (Lenoir et al., 2008; Biella et al., 2017; Freeman et al., 2018; Fazlioglu, Wan & Chen, 2020). Globally, average air temperature is predicted to increase by as much as 2.4°C–4.8°C by the end of the 21st century in the high-emissions scenario SSP5-8.5 according to the Sixth Assessment Report of the IPCC (Lee et al., 2021). Understanding the effect of increasing temperature on the survival and reproduction of plants and animals and their capacity to adapt is thus an urgent task (Aitken et al., 2008; Alberto et al., 2013).

One of the most vulnerable phases in the plant life cycle is sexual reproduction. Successful reproduction in most plant species depends on effective pollination by animals (Ollerton, Winfree & Tarrant, 2011), but also on the thermotolerance of pollen grains (He et al., 2017; Rosbakh et al., 2018), which can be highly variable in species growing in different environmental conditions (Steinacher & Wagner, 2012; Rosbakh & Poschlod, 2016). That pollen germination and pollen tube growth depend, among other factors, on temperature has been known for a long time (Brink, 1924; Das et al., 2014; Pham, Herrero & Hormaza, 2015; Shi et al., 2018; Hebbar et al., 2018). Other stages of the process of sexual reproduction are also temperature-dependent (Hedhly, Hormaza & Herrero, 2009; Zinn, Tunc-Ozdemir & Harper, 2010; Lohani, Singh & Bhalla, 2020), but pollen germination seem to be particularly sensitive to heat stress (Young, Wilen & Bonham-Smith, 2004; Zinn, Tunc-Ozdemir & Harper, 2010). Optimum temperature allows higher pollen germination success and faster pollen tube growth (Hedhly, Hormaza & Herrero, 2005), but when the temperature exceeds a species-specific optimum, pollen germination rapidly decreases and pollen tube growth stops (Lewis, 1942; Kakani et al., 2005; Zinn, Tunc-Ozdemir & Harper, 2010). Therefore, both pollen germination rate and pollen tube length can provide key information on the effects of thermal stress on the male fitness and reproduction in plants.

Global warming may thus represent a serious threat to pollen germination and the overall plant reproductive system (Hedhly, Hormaza & Herrero, 2005). Apart from increasing mean temperature, temperature extremes and heat waves are predicted to be more intense, more frequent, and to last longer (IPCC, 2021). However, the susceptibility of key stages of the sexual reproduction of plants to high temperature has been so far studied almost exclusively in crops or other commercially exploited plants (Wang et al., 2019; Bheemanahalli et al., 2019; Lovane et al., 2021; Lohani, Singh & Bhalla, 2022), but not in wild plant species, with the exception of a few studies on trees (Pigott & Huntley, 1980; Flores-Rentería et al., 2018) and forbs (McKee and Richards, 1998; Rosbakh & Poschlod, 2016).

Moreover, insights about the impact of global warming on plant populations can be gained from studies of local adaptation and acclimation to temperature along environmental, e.g. elevational gradients, to which plant species respond by with various trait changes (Aitken et al., 2008; Alberto et al., 2013; Wagner et al., 2016). Some studies indicate that plants in their native habitat exhibit higher fitness compared to plants from other populations introduced to that same location. Notably, plants from larger populations demonstrate more pronounced evidence of local adaptation (Leimu & Fischer, 2008). Additionally, Lortie & Hierro (2022) provided evidence about plant adaptation to local climate based on the analysis of seed germination, seedling growth, biomass, and other measures. Also, studies of crops confirmed intraspecific variation in pollen thermal tolerance by comparing the effects of temperature on pollen thermotolerance of different cultivars (Kakani et al., 2005; Walters & Isaacs, 2023). On the other hand, Flores-Rentería et al., (2018) tested pollen thermotolerance of *Pinus edulis* across different elevations, but their prediction that pollen from sites with higher air temperature is more tolerant to high temperature was not confirmed. The possibility of local adaptation of pollen thermotolerance to temperature thus remains an open question.

To fill this knowledge gap, we tested pollen thermotolerance of a common wild plant species, *Lotus corniculatus* s.l., by measuring pollen germination and pollen tube length in response to increasing temperature in 12 populations from two different elevations. We asked the following questions:

1. Does the pollen germination rate and pollen tube length decrease with increasing temperature? We hypothesised that pollen performance is limited by the temperature of 40 °C, because this temperature exceeds the maximum summer temperatures observed in the study region.
2. Does pollen from lowland populations have higher germination rate and pollen tube length at higher temperatures compared to pollen from highland populations? We hypothesized that pollen of plants from the lowland populations has higher termotolerance at higher temperatures because of local adaptation of lowland plants to warmer climate.

## Materials & Methods

### Study sites

The study was carried out in the southern part of the Czech Republic. We conducted our experiment at six lowland sites (389 – 451 m a.s.l.) in the surroundings of the city of České Budějovice and six highland sites (841 – 1030 m a.s.l.) in the nearby Šumava Mountains (Table 1.). We did not measure climatic data in situ because it was not necessary to obtain exact measurements of microclimatic conditions for the purpose of our study. We only aimed to select a set of highland sites generally colder than selected lowland sites. Therefore, we obtained publicly available temperature data released by the Czech Hydrometeorological Institute (https://www.chmi.cz/) from the nearest meteorological stations located within several km from the sites at a similar elevation – one station in the lowland and one in the highland (Table 1). We also downloaded climate data for each site from the Worldclim database (https://www.worldclim.org/, Fick & Hijmans, 2017) based on exact GPS coordinates of the sites. We selected the average maximum temperature of the warmest months as the most biologically relevant among the climate variables in Worldclim because our sampling was done in the summer when the temperatures reach the maximum values. The temperature values for each site were extracted based on its GPS coordinates using the R package *raster* (Hijmans, 2021). The differences in maximum temperature between the lowland (mean 23.9°C) and highland sites (mean 19.6°C) were >4°C (Table 1.). This difference in elevation represents a good proxy for the effect of global warming, because it corresponds to predictions of future global warming by 2.4°C–4.8°C by the end of the 21^st^ century in the high-emissions scenario SSP5-8.5 according to the Sixth Assessment Report of the IPCC (Lee et al., 2021).

**Table 1.** The list of study sites in the lowland and the highland with temperature details. *T*_*max*_ *warmest month* represents the average highest air temperature of the warmest month at the study sites according to Worldclim, *T*_*max*_ *three days prior* represents the average of the highest temperature in the day of collection and two days before the collection. The elevation and GPS coordinates of the centre of each site are also provided.

| Site ID | Site name | Elevation (m a.s.l.) | $T_{max}$ warmest month (°C) | $T_{max}$ 3 days prior (°C) | GPS coordinates |
| --- | --- | --- | --- | --- | --- |
| Lowland 1 | Břehov | 408 | 24.1 | 22.0 | 49.0193356N, 14.3288503E |
| Lowland 2 | Haklovy Dvory | 389 | 24.0 | 22.0 | 48.9990081N, 14.3891314E |
| Lowland 3 | Křenovice | 398 | 24.0 | 22.0 | 48.9899975N, 14.3616067E |
| Lowland 4 | Branišov | 406 | 23.9 | 25.0 | 48.9782317N, 14.4154694E |
| Lowland 5 | Třebín | 414 | 23.8 | 25.0 | 48.9665339N, 14.3844175E |
| Lowland 6 | Kvítkovice | 451 | 23.6 | 25.0 | 48.9584728N, 14.3290661E |
| Highland 1 | Strážný CHKO | 940 | 19.1 | 20.1 | 48.9290808N, 13.7092800E |
| Highland 2 | Strážný | 841 | 19.6 | 20.8 | 48.9174489N, 13.7183381E |
| Highland 3 | Křišťanov | 930 | 19.6 | 22.5 | 48.9078519N, 14.0228328E |
| Highland 4 | Koryto | 870 | 20.1 | 22.5 | 48.9315508N, 13.9978347E |
| Highland 5 | Arnoštov | 890 | 19.9 | 22.5 | 48.9061311N, 14.0014181E |
| Highland 6 | Sněžná | 1030 | 19.0 | 26.0 | 48.8961061N, 13.9393994E |

### Study species

Our study species is a dicotyledonous forb *Lotus corniculatus* L. s.l. (Fabaceae). We selected this species because of its common occurrence across the Czech Republic spanning the entire climatic gradient from the warmest lowlands to the coldest mountain ranges (Chytrý et al., 2021). *L. corniculatus* has a long flowering period (April – August) and is strictly entomogamous; the most frequent pollinators are solitary bees and bumblebees (Chytrý et al., 2021). Pollen germination in *L. corniculatus* has been studied by Wagner et al. (2016), who tested its cold tolerance and found that pollen of *L. corniculatus* requires temperatures above 5°C for germination and reached >80% germination rate already at 10°C. We are not aware of published data on the effect of higher temperatures on pollen germination in this species.

### Data collection

Before the collection of *L. corniculatus* plants, we prepared 1.5 litres of BK culture medium (Brewbaker & Kwack, 1963) modified for best performance in Fabaceae according to Tushabe & Rosbakh (2021) in the laboratory. We used 50g 10% 1L sucrose, 50 mg boric acid H_3_BO_3_, 150 mg calcium nitrate Ca(NO_3_)_2_, 100 mg magnesium sulphate heptahydrate Hu_14_MgO_11_S, 50 mg potassium nitrate KNO_3_ for medium preparation. We made up the medium with 500 ml deionized water. We calibrated the medium to pH 5.5. Finally, we autoclaved the medium for 2 hours at 120°C. We did not add agar, thus the medium remained liquid, which allowed us easier manipulation.

We collected the plants during June – July 2022 in the field. In each of 12 sites, we collected 20 individuals between 10:00 and 16:00 hours, when we observed the highest activity of pollinators. We selected plants with fresh-looking flowers (without visible signs of senescence). Entire plant stems with flowers were cut and immediately placed into plastic ziplock bags with a piece of wet paper towel and stored in polystyrene cooling boxes at ambient temperature to prevent their wilting during transport to the lab.

The experiments were conducted in the laboratory using fresh pollen during the day of collection. We released the pollen from mature anthers of five flowers per individual using entomological forceps. The pollen was placed to an Eppendorf tube with liquid BK culture medium (described above). Afterwards, we put the Eppendorf tubes in plastic racks to incubators with four different temperature levels (15, 25, 30, and 40°C). These temperature levels were designed based on other pollen germination studies (Kakani et al., 2005; Kaushal et al., 2016; Mesihovic et al., 2016; Walters & Isaacs, 2023) and our own pilot experiments to cover temperatures likely within and above the optimum temperature range. In the lowland parts of the study region, summer temperatures measured 2 m above ground occasionally exceed 30°C, but never reach 40°C according to measurements of the Czech Hydrometeorological Institute (https://www.chmi.cz/), although the temperatures of microhabitats close to the ground where the flowers are located may be higher (Scherrer & Koerner, 2009). Five replicates for each combination of site and temperature level were incubated for 20 hours in the darkness. After the incubation, the samples were placed to the freezer (-20 °C), which allows long-term storage of the samples without damage to the pollen grains (Du et al., 2019). No damage to pollen grains was apparent after taking the samples out of the freezer after approximately two months. We processed the samples (counting of pollen grains and measurements of pollen tube length) during the autumn of the same year.

In the day of pollen processing, samples were left to defrost for 15 minutes at room temperature. Using a pipette, we removed three drops (1 drop = 40 µl) of the medium. We mixed the sample content with pipette suction. Medium with pollen was retrieved with clipped pipette tips with increased diameter to avoid possible damage to the germinated pollen tubes. We put these three drops on one slide and covered them by three cover slides. We put the slides with the medium under the microscope with an attached camera (Canon EOS 77D). We observed the slides under 160x magnification and took three photos at random locations within each drop; i.e., nine photos per sample in total.

Photos obtained by a camera attached to the microscope, examples obtained under different temperatures in Figure 1, were processed in the Fiji distribution of ImageJ software (Schindelin et al., 2012). We visually counted all pollen grains in each photo and distinguished germinated pollen grains (those with a pollen tube at least the length of the pollen grain diameter). We calculated germination rate as the proportion of germinated grains divided by all pollen grains in a sample (pooled numbers from the nine photos per sample). We also measured the pollen tube length of germinated pollen grains by drawing a line along the full length of each pollen tube and measuring its length in ImageJ. Pollen tubes were measured only if they were longer than the pollen grain diameter. Burst pollen grains were not counted as germinated pollen. In total, we counted 25,427 pollen grains, mean number of pollen grains per photo was 14.5.

**Figure 1.**
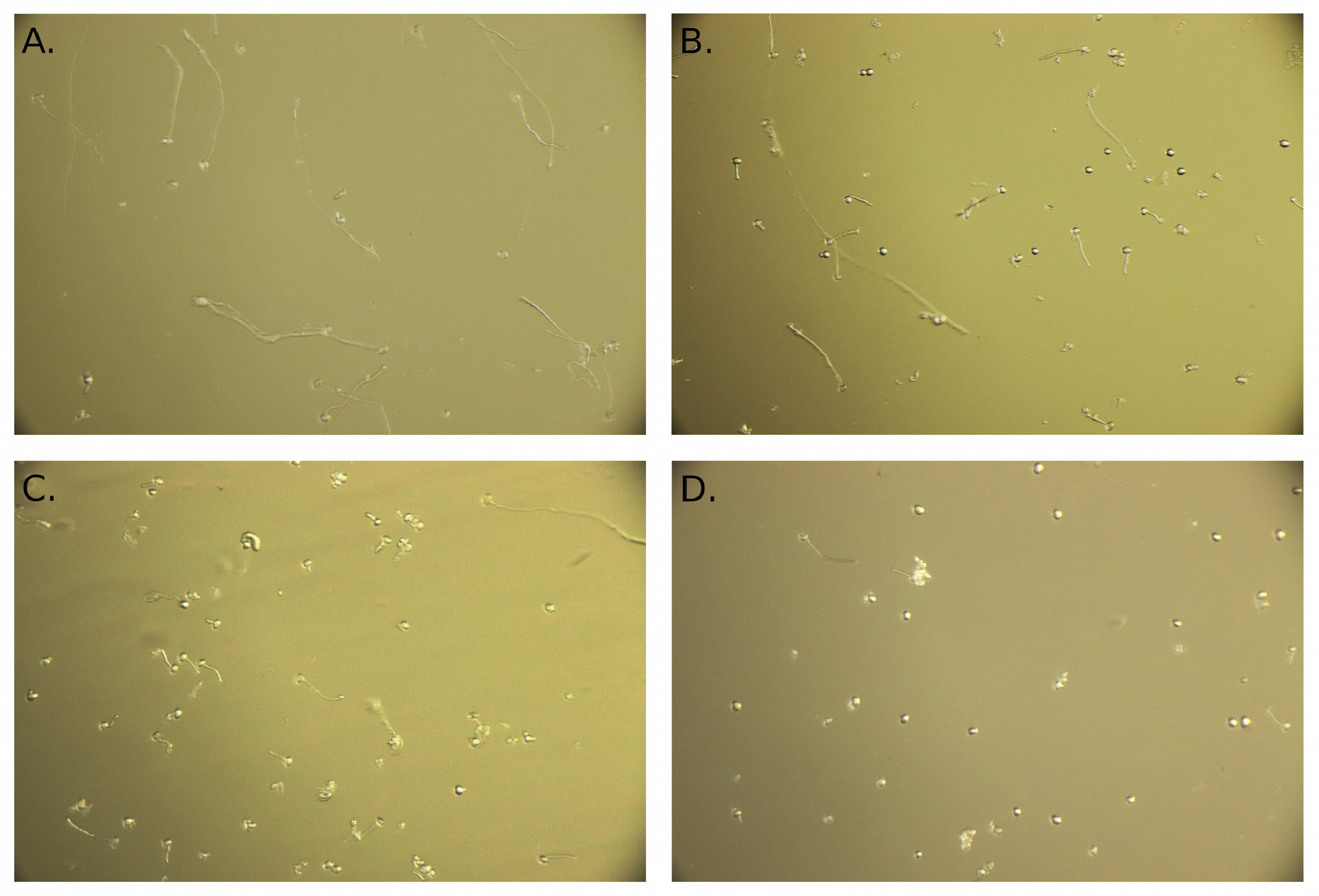
Example of photos of pollen grains. Pollen grains incubated at the temperatures of 15°C (A.), 25°C (B.), 30°C (C.), and 40°C (D.).

### Data analysis

Data from the experiment were analyzed using a generalized linear mixed model (GLMM) with the glmer function implemented in the lme4 package (Bates et al., 2015) in R version 4.2.2 (R Core Team, 2022). Temperature data from Worldclim for focal sites were extracted in R using the library raster (Hijmans, 2021).

Temperature (a factor with four levels), elevation (a factor with two levels), and their interaction were used as predictors in the analyses. Pollen germination rate (proportion of pollen grains which germinated per sample) and pollen tube length were used as response variables in two separate models (Table 2.). For pollen germination data, we used binomial error distribution and included the number of pollen grains per sample as weights. For pollen tube length data, we used gamma error distribution with log link function. We performed model diagnostics to ensure that the model assumptions were satisfied. The site of plant origin was included as a random effect in both cases. We used likelihood ratio tests to evaluate the statistical significance of the predictors. In addition, we conducted post-hoc multiple comparison tests to compare pollen germination rate and pollen tube length between all combinations of temperature levels, separately for lowland and highland plants, using the function glht in the multcomp library for R (Hothorn, Bretz & Westfall, 2008).(Hothorn, Bretz & Westfall, 2008).

**Table 2.** The results of generalized linear mixed effects models (GLMM) testing the dependence of pollen germination rate and pollen tube length on different temperature levels and elevation. Parameter estimates, their standard errors (SE), z or t values, and P values are provided for fixed effects of the elevation (lowland x highland), temperature (4 levels), and their interaction. Values of parameter estimates are provided at the scale of the linear predictors. Logit link function was used in the binomial GLMM testing the effect of elevation and temperature on pollen germination rate and log link function was used in the GLMM testing the effect of the predictors on pollen tube length. P values <0.05 are highlighted in bold.

| Predictors (fixed effects) | Pollen germination rate |  |  |  | Pollen tube length |  |  |  |
| --- | --- | --- | --- | --- | --- | --- | --- | --- |
|  | Estimate | SE | z | P | Estimate | SE | t | P |
| Intercept | -0.018 | 0.098 | -0.19 | 0.8524 | -1.305 | 0.071 | -18.38 | **< 2\*10<sup>-16</sup>** |
| Temperature 25 | -0.053 | 0.065 | -0.82 | 0.4118 | -0.136 | 0.039 | -3.49 | **0.0005** |
| Temperature 30 | -0.081 | 0.063 | -1.30 | 0.1936 | -0.425 | 0.038 | -11.18 | **< 2\*10<sup>-16</sup>** |
| Temperature 40 | -0.539 | 0.06 | -8.92 | **< 2\*10<sup>-16</sup>** | -0.546 | 0.039 | -14.17 | **< 2\*10<sup>-16</sup>** |
| Elevation Highland | 0.307 | 0.114 | 2.70 | **0.0069** | -0.289 | 0.078 | -3.71 | **0.0002** |
| Temperature 25 : Elevation Highland | -0.086 | 0.090 | -0.96 | 0.3390 | 0.026 | 0.051 | 0.50 | 0.6154 |
| Temperature 30 : Elevation Highland | -0.369 | 0.088 | -4.19 | **2.9\*10<sup>-5</sup>** | 0.182 | 0.052 | 3.51 | **0.0004** |
| Temperature 40 : Elevation Highland | 0.079 | 0.082 | 0.96 | 0.3388 | 0.093 | 0.050 | 1.88 | 0.0600 |

## Results

We observed a statistically significant effect of the interaction of the temperature of incubation and the elevation of plant origin on pollen germination rate *(χ*^2^= 41.82, P = 4.376*10^-09^, Table 2). Lowland plants had stable germination rate in the temperature range from 15 to 30°C, followed by a strong reduction at 40°C (Figure 2.). While on average between 47and 50% of pollen grains germinated at 15-30°C, pollen germination rate significantly dropped to 36% at 40°C (P<1*10^-05^ based on post-hoc tests comparing pollen germination rate at 40°C compared to the lower temperatures). On the other hand, the germination rate of pollen from highland plants significantly decreased already as the temperature reached 30°C, from 57% at 15°C and 53% at 25°C to 46% at 30°C and 45% at 40°C (P<0.001 based on post-hoc tests comparing pollen germination rates at 30°C or 40°C compared to 15°C or 25°C). In addition, the overall average pollen germination rate significantly differed between lowland and highland sites; the proportion of pollen grains which germinated was on average 8% higher in plants from the highland (z = 2.70, P = 0.0069, Figure 2.).

**Figure 2.**
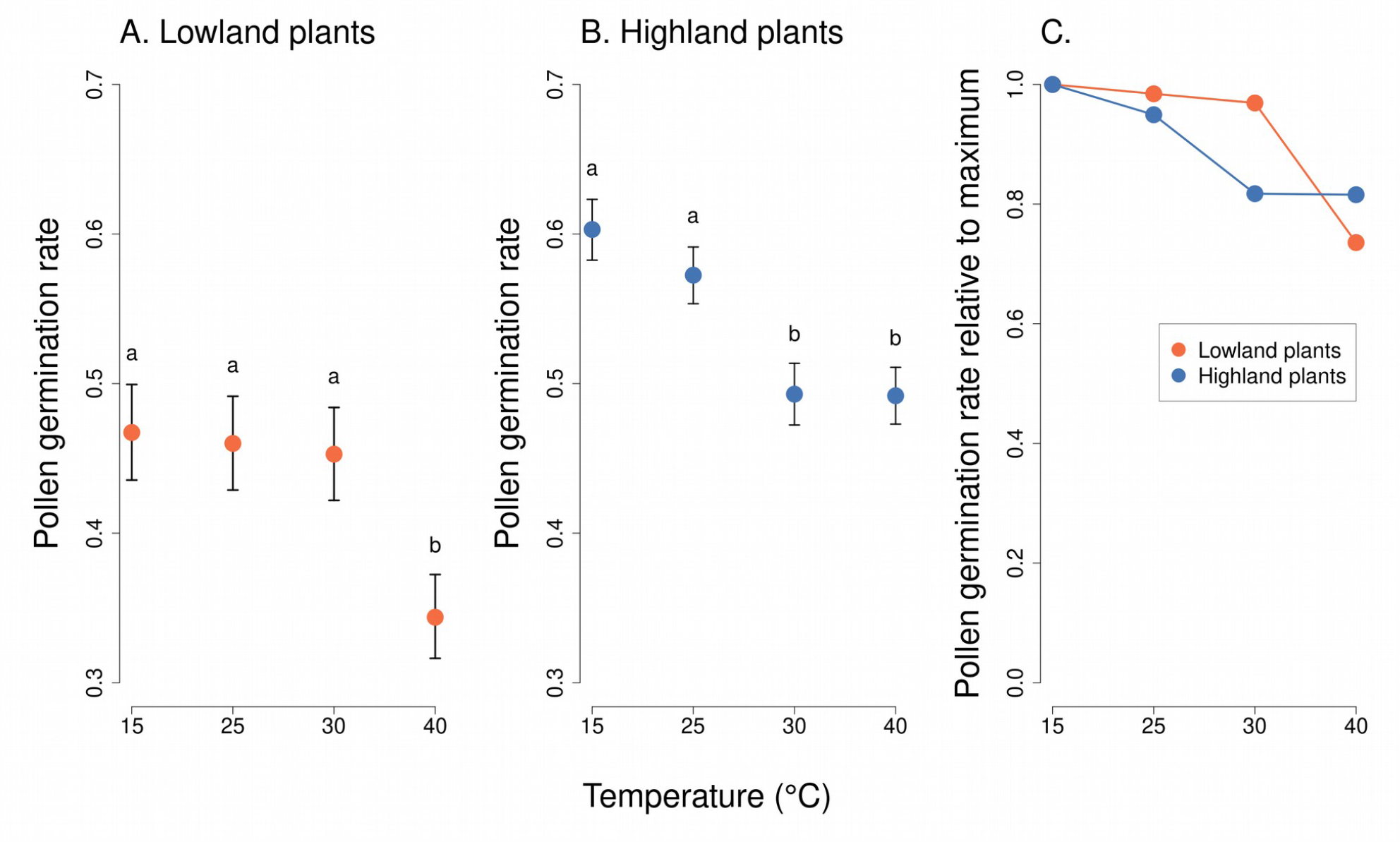
Pollen germination of plants from the lowland and the highland at four temperature levels. Mean values and SE estimated by a generalised linear mixed effects model are plotted (A. and B.). The fixed effect of temperature is displayed. Pollen germination rate is expressed as the proportion of pollen grains which germinated. The letters above the points show results of multiple comparison tests between all combinations of temperature levels, calculated separately for data on lowland and highland plants. Differences in pollen germination at temperature levels marked by a different letter were statistically significant (p<0.001 in all cases). The comparison of relative values of pollen germination rate in relation to the maximum is shown in C.

Analysis of pollen tube lengths also showed a statistically significant effect of the interaction of the incubation temperature and the elevation of plant origin (*χ*^2^ = 17.12, P = 0.0007, Table 2). However, the differences between lowland and highland sites were less pronounced than in the case of pollen germination rate (Figure 3.). Pollen tube length decreased with increasing temperature in a similar way in lowland and highland populations (Figure 3.). The average pollen tube length was slightly shorter, on average by 68 µm, in plants from the highland compared to the lowland (t = -3.709, P = 0.0002, Figure 3.).

**Figure 3.**
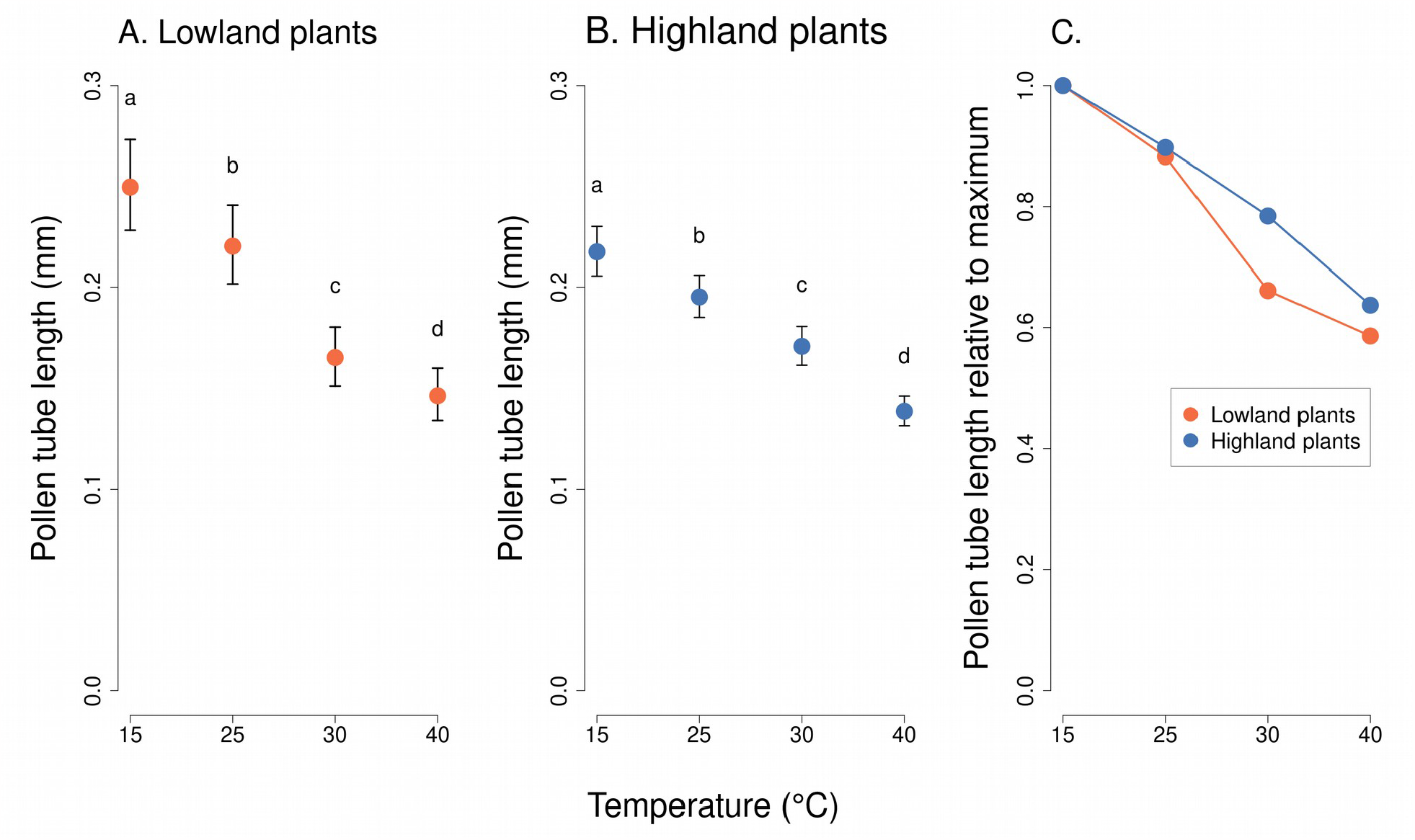
Pollen tube length of plants from the lowland and the highland at four temperature levels. Mean values and SE estimated by a generalised linear mixed effects model are plotted (A. and B.). The fixed effect of temperature is displayed. The letters above the points show results of multiple comparison tests between all combinations of temperature levels, calculated separately for data on lowland and highland plants. Differences in pollen tube length at temperature levels marked by a different letter were statistically significant (p<0.001 in all cases). The comparison of relative values of pollen germination rate in relation to the maximum is shown in C.

## Discussion

Until now, the effect of increasing temperature on plant sexual reproduction has been studied mostly in crops because of concerns about the effects of climate change on food production (Hedhly, Hormaza & Herrero, 2009; Zinn, Tunc-Ozdemir & Harper, 2010) but little attention has been paid to the effects of increasing temperature on sexual reproduction of wild plants (Rosbakh et al., 2018). Although species with frequent vegetative reproduction could persist with non-viable pollen, relying on vegetative reproduction leads to the loss of genetic variability and decreases the ability to adapt to unpredictable changes in the environment (Hedhly, Hormaza & Herrero, 2009; Eckert et al., 2010). Hence, research on the direct effects of increasing temperature on sexual reproduction of wild plants is needed to evaluate the threat posed by global warming for plant diversity.

### The effects of high temperature on pollen thermotolerance

Pollen germination rate and pollen tube length of *Lotus corniculatus* significantly decreased at high temperature. The pollen germination rate was reduced at 40°C in the lowland population and also at 30°C in the highland population, compared to lower temperatures. Although very rarely studied in wild plant species, a negative effect of high temperature on pollen germination rate has been observed in many species of crops (Zinn, Tunc-Ozdemir & Harper, 2010; Kaushal et al., 2016). For example, pollen germination rate of rice *(Oryza sativa)* significantly decreased when exposed to temperature higher than 32 – 35 °C in a study of Rang et al. (2011) and 38°C in a study by Shi et al. (2018). Also, almond *(Prunus dulcis)* had reduced pollen performance at 35°C, which inhibited pollen germination (Sorkheh et al., 2011), similarly to spring wheat *(Triticum aestivum* L.) exposed to 34°C (Bheemanahalli et al., 2019). Optimum temperature for pollen germination in cotton *(Gossypium hirsutum)* was approximately 30°C with a sharp decrease at higher as well as lower temperatures (Kakani et al., 2005). Among crops from temperate regions, pollen germination rate of the northern highbush blueberry *(Vaccinium corymbosum)* was reduced at temperatures exceeding 35°C and almost completely prevented at 40° (Walters & Isaacs, 2023).

Pollen tube length was also significantly reduced with increasing temperature, but in this case, the reduction was more gradual. Previous studies of crops also provide examples of a negative effect of high temperatures on pollen tube growth. For instance, Longan *(Dimocarpus longan)*, a subtropical fruit tree, showed decreased pollen tube length at 40 °C (Pham, Herrero & Hormaza, 2015; Pham et al., 2015). Also. Hebbar et al. (2018) found out that some cultivars of coconut (*Cocos nucifera)* have shorter pollen tubes at temperatures around 40 °C. Decreasing pollen tube length with increasing temperature was also observed in *Pistacia* spp (Acar & Kakani, 2010), in cotton *(Gossypium hirsutum)* at temperatures above the optimum temperature of 30°C (Kakani et al., 2005), and in the northern highbush blueberry *(Vaccinium corymbosum)* also at temperatures exceeding 30°C (Walters & Isaacs, 2023).

### Signs of local adaptation

As hypothesized, we observed differences in the effect of increasing temperature on the pollen germination rate of lowland and highland populations of *L. corniculatus*. While pollen germination rate was stable at the incubation temperature of 15-30°C in the lowland plants and a reduction of pollen germination rate was observed only at 40°C, pollen germination rate of the highland plants was reduced already when the temperature of incubation reached 30°C. We suggest that this is due to local adaptation of the lowland plant populations to higher temperatures. The difference in the elevation of the lowland and highland sites was over 400 m and climate data show that the differences in long-term average temperatures between the lowland and highland sites exceed 4°C (Table 1). Plants in the lowland populations thus experience substantially higher temperatures during their flowering period and this provides selection pressure to adapt to higher temperatures.

Variation of plant traits along environmental gradients, which suggests local adaptation of plant populations to varying conditions has been observed in many species (Aitken et al., 2008; Alberto et al., 2013; Wagner et al., 2016). A large amount of indirect evidence comes from numerous common garden experiments which showed that various plant traits and different measures of plant performance depend on the origin of populations along the environmental gradient. Direct evidence from reciprocal transplant experiments is mixed, but some studies showed that plants at their native site had higher fitness than plants from other populations introduced to that site, with plants from larger populations showing stronger evidence for local adaptation (Leimu & Fischer, 2008). A recent meta-analysis of seed and seedling transplant experiments provided a strong overall support for plant adaptation to local climate based on the analysis of seed germination, seedling growth, biomass, and other measures (Lortie & Hierro, 2022). However, the only reciprocal transplant study on *Lotus corniculatus* we are aware of found no evidence for adaptation to climate based on comparisons of total biomass and fecundity of plants reciprocally transplanted among three sites (Macel et al., 2007). Despite the large body of research on local adaptation, we are not aware of previous studies focusing on adaptation of pollen thermotolerance to increasing temperature, with the exception of a study of *Pinus edulis* along an elevational gradient (Flores-Rentería et al., 2018). However, that study did not confirm the hypothesis that pollen from plants growing at lower elevations is more tolerant to higher temperatures.

Based on the design of our experiment, we could not obtain a precise estimate of the optimum and maximum temperature for pollen germination of *L. corniculatus*. However, our choice of temperature levels reflected the local climate. Specifically, we observed that pollen germination rate of highland plants was significantly reduced at 30°C compared to 15 and 25°C, while lowland plants had stable pollen germination rate at 15, 25, and 30°C and pollen germination rate was significantly reduced only at 40°C. Data from nearby meteorological stations show that while maximum daily temperatures in the lowland (ca. 400 m a.s.l.) regularly exceed 30°C, they very rarely do so in the highland (ca. 900 m a.s.l.). For example, in 2022, 30°C was exceeded on 15 days at a station in České Budějovice (elevation 381 m a.s.l.) near the lowland sites, but never at a station in Nicov (elevation 935 m a.s.l.) near the highland sites, where the maximum recorded temperature was 29.0°C. This is reflected in the different effects of increasing temperature on the germination rate of pollen grains of plants from the highland compared to the lowland populations.

### Possible acclimation and confounding factors

A caveat in our interpretation of the differences between the lowland and highland populations is that pollen thermotolerance may be affected also by the temperature the plants experienced in the field prior to the collection of pollen; i.e., there may be an effect of acclimation leading to acquired tolerance of high temperature (Sung et al., 2003). Due to acclimation, plants are able to express heat stress-responsive genes (Müller & Rieu, 2016) and also photosynthesize more efficiently in new climate conditions (Yamori, Hikosaka & Way, 2014). A study by Firon et al. (2012) also demonstrated a positive effect of previous exposure of tomato plants to high non-lethal temperature on their pollen thermotolerance under subsequent heat stress.

Accounting for the effect of acclimation in plants collected in the field is not trivial. To evaluate whether acclimation might bias our results, we compared temperature data from the meteorological stations near the lowland and highland sites. We specifically looked up the maximum air temperature on the day of sample collection and two days before that and compared the average values at individual sites. The average maximum air temperature during this time period ranged from 22.0 to 25.0°C at the lowland sites and between 20.1 and 26.0°C at the highland sites (Table 1). Plants in the lowland and the highland thus experienced very similar temperatures prior to pollen collection and we argue that the difference in the effect of high temperature on pollen germination in the lowland and highland plans observed in the lab is thus unlikely to be biased by the effect of differences in temperature experienced by the plants in the field.

Apart from the effect of temperature, we observed that the average pollen germination rate was higher in plants from the highland populations compared to the lowland populations, but the pollen tubes were slightly but statistically significantly longer in plants from the lowland populations. We do not have a good explanation for these differences, although we suspect that higher pollen germination rate in plants from the highland might be related either to differences in general plant condition possibly caused by higher water availability in the mountains (Chytrý, 2017), or by differences in air humidity, which is known to affect pollen thermotolerance (Aronne, 1999; Leech, Simpson & Whitehouse, 2002). The negative effect of drought on pollen germination was confirmed in multiple crops, e.g. in maize *(Zea mays* L.) (Bheemanahalli et al., 2022), soybean *(Glycine max)* (Poudel et al., 2023), and rice *(Oryza sativa)* (Rao et al., 2019). We did not measure precipitation at the sites, but data from a drought monitoring project by the Global Change Research Institute of the Czech Academy of Sciences (https://www.intersucho.cz/) show that the area where the highland sites were located consistently has higher soil water content than the lowland area, including during the time of our sampling. Another important factor could be whether the plant is exposed to direct sunshine most of the day or is located in the shadow (Corbet, 1990), which we did not record.

Interestingly, a previous experiment on pollen germination of *Pinus edulis* showed variable pollen germination across an elevation gradient, with the highest pollen germination rate at intermediate elevations where the plants are in their optimal conditions (Flores-Rentería et al., 2018). Also, Rosbakh & Poschlod (2016) observed variable pollen germination rate of multiple forb species across an elevational gradient. They confirmed that pollen of lowland species germinates and grows pollen tubes at relatively high temperatures compared to highland species which perform better at lower temperatures. In addition, McKee & Richards (1998) showed that increasing temperature may hamper seed set of several *Primula* species. On the other hand, low temperature can also negatively affect pollen performance as in the case of *Tilia cordata* where it prevents sexual reproduction at the northern distribution limit (Pigott & Huntley, 1980). Minimum temperature requirements for pollen germination and pollen tube grows are also species specific depending on their elevation of occurrence (Rosbakh & Poschlod, 2016) and time of the year when they are flowering (Wagner eta al. 2016).

Multiple studies demonstrated that different environmental drivers, such as temperature, water, and nutrient availability often have interactive effects on plant vegetative growth, floral traits, and interactions with pollinators (Hoover et al., 2012; Descamps et al., 2018; Akter & Klečka, 2022). Hence, understanding the effects of a broader set of local environmental conditions on pollen thermotolerance will certainly be an interesting avenue for future research.

## Conclusions

Our experiment with pollen germination in lowland and highland populations of *Lotus corniculatus*, which were tested across four temperature levels, provides evidence that pollen germination is more sensitive to high temperatures in highland plants compared to lowland plants. This is likely the result of local adaptation to higher temperatures in lowland populations. In addition, we found a negative effect of increasing temperature on pollen tube length, which may further prevent fertilization at high temperatures. We conclude that more research on the temperature-dependence of pollen thermotolerance of wild plants is needed to understand the impact of climate change on plant reproduction. Our data show that increasing temperature may hamper sexual reproduction of wild plants, similarly to crops, but differences among populations suggest potential for adaptation.

## Acknowledgements

We would like to thank Jenna Walters for discussions about pollen germination, Dagmar Hucková and Davide Panzeri for their help with the preparation of culture medium, and Pavel Duda for his help in the field. We would also like to thank Nicola Tommasi and Zdeněk Faltýnek Fric for useful comments about data analyses. Last but not least, we are grateful to Eva Kaštovská and Petr Čapek for providing their incubators for the experiment.

